# Scattering-Based Super-Resolution Optical Fluctuation Imaging

**DOI:** 10.1101/2024.09.02.610873

**Authors:** Shimon Yudovich, Gregor Posnjak, Lior Shani, Eti Teblum, Tim Liedl, Jörg Enderlein, Shimon Weiss

## Abstract

Super-resolution optical imaging has become a prominent tool in life and material sciences, allowing one to decipher structures at increasingly greater spatial detail. Among the utilized techniques in this field, super-resolution optical fluctuation imaging (SOFI) has proved to be a valuable approach. A major advantage of SOFI is its less restrictive requirements for generating super-resolved images of neighboring nanostructures or molecules, as it only assumes that the detected fluctuating light from neighboring emitters is statistically uncorrelated, but not necessarily separated in time. While most optical super-resolution microscopies depend on signals obtained from fluorescence, they are limited by photobleaching and phototoxicity. An alternative source for optical signals can be acquired by detecting the light scattered from molecules or nanoparticles. However, the application of coherent scattering-based imaging modalities for super-resolution imaging has been considerably limited compared to fluorescence-based modalities. Here, we develop scattering-based super-resolution optical fluctuation imaging (sSOFI), where we utilize the rotation of anisotropic particles as a source of fluctuating optical signals. We discuss the differences in the application of SOFI algorithms for coherent and incoherent imaging modalities, and utilize interference microscopy to demonstrate super-resolution imaging of rotating nanoparticle dimers. We present a theoretical analysis of the relevant model systems, and discuss the possible effects of cusp artifacts and electrodynamic coupling between nearby nano-scatterers. Finally, we apply sSOFI as a label-free novelty filter that highlights regions with higher activity of biomolecules and demonstrate its use by imaging membrane protrusions of live cells. Overall, the development of optical super-resolution approaches for coherent scattering-based imaging modalities, as described here, could potentially allow for the investigation of biological processes at temporal resolutions and acquisition durations previously inaccessible in fluorescence-based imaging.

## 1. Introduction

Super-resolution optical microscopy has become an important tool in modern biological research, material science, and nanoscience. Advances in super-resolution imaging such as stimulated emission depletion (STED) microscopy [1], photo-activated localization microscopy (PALM) [2], stochastic optical reconstruction microscopy (STORM) [3], or structured illumination microscopy (SIM) [4] and their derivatives reached optical resolutions of a few tens of nanometers, and more recently, even down to single nanometers [5]. This dramatic resolution enhancement has led to several significant new discoveries [6–8]. Most of the existing methods rely on fluorescence that, while guaranteeing high specificity and sensitivity, is associated with significant shortcomings, most notably - photobleaching, phototoxicity, and/or the requirement of non-native gene expression. An alternative to fluorescence-based imaging can be found in imaging modalities that utilize elastic scattering from molecules or particles. Such imaging modalities, such as dark-field microscopy [9], reflection interference contrast microscopy (RICM) [10], and interference scattering microscopy (iSCAT) [11, 12], have been developed to allow for sensitive optical imaging without the need for fluorescence labeling.

In interference-based microscopes such as RICM and iSCAT, the sample is illuminated by incident laser light, and one records the interference between the scattered electric field and a fraction of the back-reflected incident light. The interference between the weak scattered light with the orders-of-magnitude stronger incident light leads to an amplification of the weak scattering signal by orders of magnitude (heterodyne gain); this amplified signal is linearly proportional to the scattered electric field amplitude and is thus proportional to the third power of the scatterer’s size. This makes interference microscopy orders of magnitude more sensitive than dark-field imaging. The combination of single-molecule sensitivity, the possibility of label-free imaging, and the absence of photobleaching, gives interference microscopy enormous potential and enables temporally unlimited, label-free imaging with maximal sensitivity. Label-free scattering-based microscopy in its different forms has already proven to be relevant in the study of actin networks [13], microtubules in mitotic spindles [14] and surface assays [15], mitosis characterization [16], initiation and dynamics of liquid-liquid phase separation [17], mass quantification of cells [18] and single proteins [19], neuronal deformation due to action potentials [20], and many other applications in the realms of life and material sciences [21, 22]. However, while such microscopy techniques allowed for ultra-sensitive detection of small scatterers, the achievements of interference-based super-resolution imaging have been limited compared to their fluorescence-based counterparts [23].

In this work, we investigate the application of super-resolution optical fluctuation imaging (SOFI) [24] for sub-diffraction scattering-based imaging, which we term sSOFI. SOFI, which relies on cumulant analysis of stochastic fluctuations of optical signals, is advantageous over localization-based approaches since it does not require neighboring nano-objects to emit light in a temporally separated manner. The basic working principle of SOFI can be described as follows: assuming each image is generated by *N* discrete emitters/scatterers at positions **r***_j_* (1 ≤ *j* ≤ *N)*, with the intensity at the camera plane *F*(*r*, *t*) proportional to the sum of each individual time-dependent (fluctuating) emitted/scattered intensity:

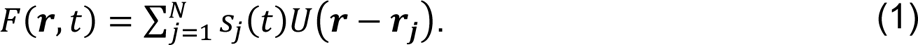

Here, *U*(*r* − *r*_*j*_) is the point spread function (PSF) of the applied imaging modality and *s*_*j*_(*t*) is the time-dependent intensity of the *j*^th^ emitter/scatterer. The above assumption, that the formed image is composed from the addition of independent and non-correlated PSFs from different point-particles, is not valid for dark-field imaging, as further discussed in section 3.A. In second-order SOFI, one calculates the second-order cumulant image *C*_2_(*r*, *τ*), which is defined by

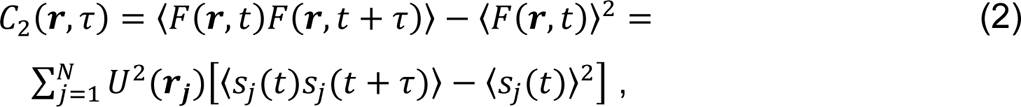

where the angular brackets denote averaging over time *t*, and *τ* is a freely chosen correlation time. The last expression on the right-hand-side is the sum of second-order cumulants of the optical signal trajectory *s*_*j*_(*t*), assuming all cross-cumulant contributions between different emitters/scatterers are negligible. It is important to note that this assumption requires statistical independence of amplitude fluctuations between different emitters/scatterers. Then, equation (2) shows that the second-order SOFI image is formed with the square of the original PSF, resulting in a resolution enhancement by a square-root of two compared to the pure intensity image. Theoretically, the size of the PSF for an *n*^th^-order SOFI image is reduced by a factor of *n*^1/2^, as compared to the original acquisition. Applying deconvolution (Fourier reweighting) [25, 26] in conjunction with knowledge of the system’s optical PSF can provide an additional enhancement of *n*^1/2^, bringing the total theoretical resolution enhancement factor to *n*. The fluctuations can originate from fluorescent proteins [27], organic dyes [28], quantum dots [24], or carbon nanodots [29]. Other types of optical fluctuations, such as those originating from diffusion [30], diffusion-controlled FRET [31], or stochastic speckle illumination [32], have also been exploited for SOFI. To date, SOFI has been implemented through a number of imaging platforms including wide-field microscopy [33], total internal reflection fluorescence microscopy [27], multi-plane wide-field fluorescence microscopy [34], spinning-disk confocal microscopy [35], and light sheet microscopy [36]. In addition, dynamic speckle illumination has been used to extend SOFI to acousto-optic and photo-acoustic tomography [37]. The major advantages of SOFI include simplicity, low cost, compatibility with different imaging platforms, the availability of a wide variety of blinking probes, flexibility in imaging conditions [38], low excitation power ideally suited for imaging living cells, and a useful trade-off between spatial and temporal resolutions.

In this work, we show that SOFI analysis could be applied for interference microscopy data. However, it is incompatible with dark-field microscopy, as the formed image contains an additional cross-term component between two neighboring emitters that inherently fluctuates in a manner correlated with the two emitters. Next, we experimentally demonstrate this approach by performing interference imaging of gold nanorod (AuNRs) dimer structures formed by DNA origami. Such engineered nano-rulers with conjugated scattering objects at pre-defined positions provide a testbed for the development of scattering-based super-resolution. Here, polarization-sensitive imaging of AuNRs undergoing rotational diffusion produces movies with fluctuating scattered fields, analogous to fluorescence blinking, which is an essential component for applying SOFI and localization-based super-resolution routines. Finally, since the weight of each pixel in a SOFI image is produced by the cumulant of the corresponding detected intensity trajectory, it is possible to apply sSOFI as a novelty filter that highlights dynamic processes in a label-free manner. We demonstrate this approach by interference imaging of membrane protrusions in live cells that exhibit fluctuating dynamics, and show that sSOFI offers a convenient approach to analyze such movies and produce fluctuation-sensitive contrast.

## 2. Materials and Methods

### A. Gold Nanorod Monomers and Dimers

Gold nanorods were synthesized according to a modified protocol from Gonzalez-Rubio et al. [39]. For the synthesis, all glassware and stirring bars were cleaned with aqua regia. For the Au seed growth, we mixed 5 mL 0.5 mM HAuCl_4_, 5 mM 0.2 M CTAB (TCI) and 0.6 mL of freshly prepared 0.01 M NaBH_4,_ which was diluted to 1 mL prior to mixing. The mixture was vigorously stirred at 1200 RPM for about 2 min, during which the color should change from yellow to brownish yellow. At that point we stopped the stirring and aged the mixture for 30 min. The AuNR growth solution was prepared by adding 0.9 g CTAB and 0.11 g Na-CRESO to 25 mL of MilliQ water at around 60 °C in a 100 mL Erlenmeyer flask. The solution was kept on a hot plate at 30 °C and mixed at 300 RPM with care taken to avoid forming foam, before adding 0.6 mL of 4 mM AgNO_3_. The mixture was kept undisturbed at 30 °C for 15 min. Then, we added 25 mL of 1 mM HAuCl_4_ and mixed for 15 min at 400 RPM. Afterwards, we added 0.1 mL of 0.064 M ascorbic acid (freshly prepared) and mixed the solution at the maximum available rate (1200-1500 RPM) for 30 s, at which point the solution should be colorless. We used the growth solution immediately by adding 120 µL of the seed solution during mixing, mixing for 30 s, and left undisturbed for 12 h at 30 °C. To wash the AuNR, we centrifuged the mixture for 45 min at 15k RCF and 30 °C, removed the supernatant, added warm 0.1 M CTAB, resuspended the AuNR by short mixing and ultrasonification, and centrifuged again at the same conditions. After removing the supernatant, the AuNRs were transferred to a 2 mL eppendorf tube, diluted with warm 0.1 M CTAB, resuspended by short mixing and ultrasonification, and centrifuged for 15 min at 15k RCF and 30 °C. After the removal of the supernatant, 0.1 % SDS was added, and the AuNRs were resuspended and centrifuged as before. The supernatant was removed before dilution with 0.1 % SDS to OD = 60. The AuNRs were functionalized by mixing the rods (OD = 16 in 0.1 % SDS) in a 6:5 volume ratio with a mixture of thiolated T-30 and T-8 DNA strands (Eurofins, 100 µM concentration; 1:9 volume ratio) and freezing them at -80 °C for 30 min [40]. Excess DNA strands were removed by five rounds of Amicon spin filtration (100 kDa) for 6 min at 5k RCF and 25 °C prior to hybridization with the DNA origami structure.

The DNA origami structure used for the formation of the AuNR dimers was the rectangular Rothemund DNA origami structure (RRO), with eight biotinylated staple strands (Eurofins Genomic) for attachment to the BSA–biotin–streptavidin treated surface [41]. Dedicated binding strands for DNA-functionalized AuNRs and fluorophores were designed using the Picasso software package [41], with the binding strands’ positions shown in Fig. S1. Oligo strands at two opposing corners of the rectangular structure, approximately 80 nm apart, were modified on their 3’ ends with the sequence: 5’-TT CATATGAATTGCATGGTACC AAAAAAAAAAAAAAAAAAAAAAAAAAAAAA-3’, where the first TT nucleotides are a spacer between the origami and the anchor strand, and the A-30 part is the particle binding sequence that hybridized with the T-30 strands on the AuNRs. The 20 nt part between the TT and the poly-A is a spacer that was hybridized by the addition of a complementary strand with the sequence: 5’-GGTACCATGCAATTCATATG-3’. In order to fluorescently label the DNA origami structures, oligo strands in four central positions of the structure were modified on their 3’ ends with the sequence: 5’-TTCCTCTACCACCTACATCACA-3’, which was then hybridized with the complementary strand modified with an ATTO 647N fluorophore on its 5’ end (IDT). The folding mixture for forming the DNA origami structure, comprised of the single-stranded p7249 DNA scaffold (10 nM) and all staple strands (100 nM each) in buffer containing 10 mM Tris-HCl,1 mM EDTA (pH adjusted to 8.0 with NaOH) and 12.5 mM MgCl_2_, was subject to thermal annealing in a thermal cycler (Techne), in which the reaction mixture was first heated to 80 °C for 5 min. Then, gradual cooling from 60 to 10 °C was applied, with a 1°C decrease every 5 min. Excess DNA strands were removed by five rounds of Amicon spin filtration (100 kDa) for 15 min at 2k RCF and 4 °C. The folded and purified DNA origami structures were then immobilized on glass coverslips precoated with BSA–biotin– streptavidin. For coating, coverslips were first cleaned by 5 min sonication in detergent solution (2% Hellmanex III; Hellma), double-distilled water, and acetone, then treated with 1 mg/ml BSA-biotin (Sigma-Aldrich) for 15 min and 0.5 mg/ml streptavidin (ThermoFisher Scientific) for 5 min, both in 10 mM Tris-HCl, 100 mM NaCl and 0.05 % (vol/vol) Tween 20 (pH 8). Multiple rounds of buffer exchange between and after coating steps were performed with the same buffer. DNA origami structures were diluted in 5 mM Tris-HCl, 10 mM MgCl_2_, 1 mM EDTA, and 0.05 % (vol/vol) Tween 20 (pH 8) and introduced to the coverslip until sufficient surface covering was achieved. After multiple rounds of buffer exchange, AuNRs at a saturating binding concentration, diluted in the same buffer as the DNA origami, were introduced to the coverslip and allowed to bind to the DNA origami construct for 30 min, followed by extensive buffer exchanges.

AFM measurements were conducted using a scanning probe microscope (Bio FastScan; Bruker, Karlsruhe, Germany). Images were obtained with a silicon probe (Fast Scan B, Bruker) in a soft tapping mode with a spring constant of 1.8 N/m. The cantilever was operated at a resonance frequency of approximately 450 kHz in an air environment. Image acquisition was performed in the retrace direction at a scanning speed of 1.6 Hz and 512 samples per line resolution. Image processing and artifact corrections were performed using an open-source software package (Gwyddion) [42].

### B. Interference Scattering Microscopy

Interference scattering and fluorescence microscopy were performed on a home-built imaging system, as illustrated in Fig. S2. Briefly, 470 nm and 640 diode lasers (LDH D-C-470 and LDH-D-C-640, PicoQuant), driven by picosecond laser driver (PDL 800-D, PicoQuant), were coupled into a single-mode fiber (P3-405BPM-FC-2, Thorlabs). All imaging was performed using the laser drivers’ pulsed mode, as it allowed for a shorter coherence length, which is advantageous for eliminating interference patterns created due to back-reflections in the optical path. The incoming laser beam was collimated out of the fiber using a 10x objective (PLN10X, Olympus), linearly polarized (LPVISE100-A, Thorlabs), and weakly focused (LA1908-A, Thorlabs) onto the back focal plane of a 150x oil-immersion objective (UAPON150XOTIRF, Olympus). The back-reflected and scattered field and the emitted fluorescence were split by a multi-band dichroic mirror (89402bs, Chroma) and imaged onto two different cameras. The back-reflected and scattered fields were reflected with a wedged beamsplitter (BST16, Thorlabs) and focused (AC508-500-A, Thorlabs) onto an sCMOS camera (ORCA-Flash4.0 V3, Hamamatsu), resulting in an effective pixel size of ∼14.5 nm. The emitted fluorescence was focused (AC508-180-A, Thorlabs) onto a second sCMOS camera (Prime BSI, Teledyne Photometrics), resulting in an effective pixel size of ∼45.8 nm. The fluorescence images shown in Fig. S7 were captured with 2 x 2 pixel binning. All images, apart from the membrane protrusions shown in Fig. 7, were acquired using a 640 nm laser, close to the longitudinal plasmonic resonance of the AuNRs used in this work. Fig. 6 was acquired using a 470 nm laser, with both the interference and fluorescence channels imaged onto the same sCMOS camera, adding a longpass filter (FEL0600, Thorlabs) for the detection of the fluorescence image. The sample was held on a three-dimensional piezo stage (P-562.3CD, Physik Instrumente). In order to remove the speckle background for all imaging performed on AuNRs, average background images were produced by acquiring movies composed of 1000 frames captured while oscillating the stage parallel to the sample plane. Background correction was performed by dividing raw images with the averaged background images [43], and was applied to all imaging of AuNRs monomers and dimers. However, background correction was not applied to live cell imaging since the imaged region was too crowded to allow the acquisition of background resulting solely from the glass interface. Custom-written LabView (National Instruments) and MATLAB (MathWorks) software were used for all instrument control. Fluorescent beads (Spherotech) were sparsely dispersed on a coverslip and imaged in order to align the two cameras with respect to each other.

### C. Colocalization with Scanning Electron Microscopy

In order to compare, correlate, and benchmark sSOFI results, specific regions of interest were localized by scanning electron microscopy (SEM) after optical imaging. For this purpose, we fabricated indexed chrome micro-grids on top of glass coverslips (Fig. S3). Each grid comprised an array of 20 µm x 20 µm square cells fabricated via a standard photolithography process. Briefly, PMGI SF5 and AZ 1518 photoresist (MicroChemicals) were spin-coated on clean glass coverslips (Paul Marienfeld GmbH). The substrate was exposed to the grid pattern via a maskless lithography system (MLA 150, Heidelberg Instruments). Then, a 5 nm chrome layer was sputtered (Bestec GmbH) on the developed substrate. Finally, substrates were incubated in NMP for photoresist removal. Following optical imaging, the sample was allowed to dry overnight and then sputtered with an iridium layer. SEM imaging of AuNRs was performed on a high-resolution scanning electron microscope (Magellan 400L, ThermoFisher), with the optically imaged regions of interest located using the indexed micro-grid.

### D. Labeling and Imaging of Membrane Protrusions

For interference and fluorescence imaging of membrane protrusions, human embryonic kidney cells (HEK293) were allowed to adhere to glass coverslips, followed by membrane labeling. Briefly, HEK293 cells were cultured in Dulbecco’s modified eagle medium containing 10 % fetal calf serum, 2 mM glutamine, 100 units/mL penicillin, and 100 µg/mL streptomycin and grown in 5 % CO_2_ at 37 °C. Cells were seeded on glass coverslips precoated with poly-D-lysine (Sigma-Aldrich). After plating, cells were allowed to adhere overnight and then labeled with 10 µM di-8-ANEPPS (Biotium) in a buffer solution (140 mM NaCl, 2.8 mM KCl, 2 mM CaCl2, 2 mM MgCl2, 10 mM glucose, and 10 mM HEPES, pH 7.4) for 10 minutes at room temperature. Following labeling, cells were washed twice with the same solution, which was also used as an imaging solution. All imaging was performed at room temperature shortly after membrane labeling.

## 3. Results and Discussion

### A. Theoretical Analysis of Scattering-based SOFI

In order to study the application of SOFI algorithms to scattering-based imaging modalities and compare them with their use for conventional fluorescence-based imaging, simulated movies of neighboring emitters/scatterers were produced. Assuming two non-interacting fluorescent emitters or elastic scatterers that form an image in an incoherent or coherent superposition, respectively, each forming a PSF approximated by an Airy disk. In order to introduce a source temporal intensity fluctuation, we modeled each emitter/scatterer as an anisotropic emitter/scatterer rotationally diffusing in an isotropic manner, with the detected field from each particle calculated as the projection on an axis that is orthogonal to the sample plane. Rotational diffusion was simulated by implementing Monte Carlo simulations of a random walk on a sphere according to the algorithm described in Ref. [44]. All simulations and analyses were performed with MATLAB (MathWorks). Simulated frames were produced by assuming a coherent/incoherent superposition of the fields described by each PSF weighted by the corresponding time-varying amplitude. Incoherent images were produced by adding the intensities of either contributing PSF at each simulated pixel position. In reflection interference microscopies such as RICM and iSCAT, one captures the interference pattern between the back-reflection of the incoming beam from the glass-water interface and the light scattered by the object of interest. In the case of two point scatterers, the (time-dependent) intensity distribution *I*(*r*, *t*) of this image at the camera plane can be described by:

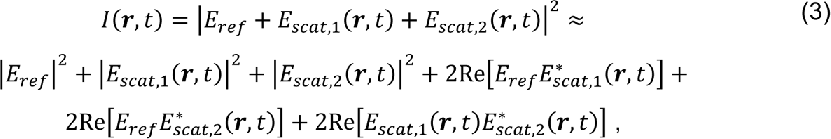

where *E*_*ref*_ and *E*_*scat*,1/2_ are the electric field amplitude of the back-reflected (reference) and scattered light from the 1^st^ and 2^nd^ particle, respectively. An important precept in interference microscopies is that the intensity of the back-reflected light is orders of magnitudes stronger than that of the light scattered by sub-micron objects, so that the term proportional to 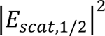 can be neglected, while the mixed-term 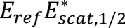 amplifies the scattered field considerably. In contrast, in the case of no reference field *E*_*ref*_ = 0, where only the scattered light is detected at the camera plane, as in dark-field imaging, the cross term between *E*_*scat*,1_ and *E*_*scat*,2_ becomes a significant contribution to the formed image.

Fig. 1 shows SOFI auto-cumulant images (*τ* = 0) up to the 8^th^-order calculated from simulated movies of two such neighboring point sources, each undergoing stochastically independent rotational diffusion, assuming four different imaging modalities: (i) incoherent (fluorescence-like) imaging, (ii) dark-field imaging (*E*_*ref*_ = 0), (iii) interference imaging with the reference field amplitude equal to the maximum scattered amplitude of each particle (*E*_*ref*_ = *E*_*scat*_), and (iv) interference imaging with the reference field is x100 greater than the maximum scattered amplitude of each particle (*E*_*ref*_ = 100*E*_*scat*_). Several observations can be deduced from the results shown in Fig. 1. First, both dark-field imaging (Fig. 1ii), and interference-based imaging with a reference field that is not significantly more intense than the scattered fields (Fig. 1iii), contain cross-terms between the two scatterers in the image plane. Such cross-terms can be viewed as artificial objects introduced to the image plane, with fluctuation dynamics that are correlated with the fluctuation dynamics of both scatterers. Since the principal assumption in SOFI algorithms is that the time-varying amplitude of each contributing PSF is statistically independent from its neighboring scatterer, these two regimes are not suitable for the formation of super-resolved images with SOFI algorithms.

**Fig. 1.**
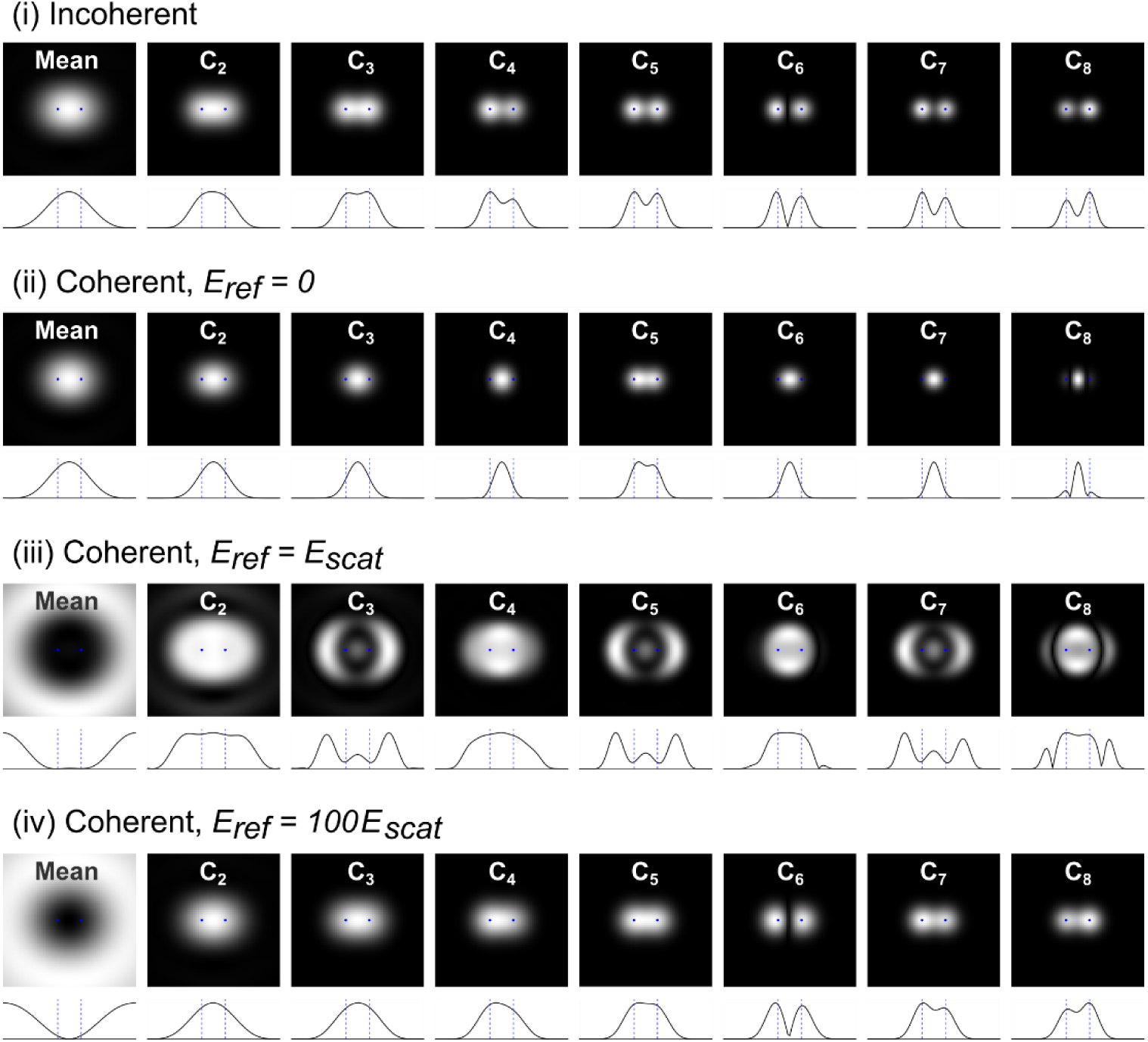
SOFI analysis of simulated movies of fluctuating sources imaged by different modalities. Simulated movies of two neighboring rotationally diffusing emitters were produced by assuming each emitter forms an independent PSF described by an Airy function, with a time-varying amplitude fluctuating as a random walk on a sphere projected into one of its axes. For identical amplitude fluctuation trajectories, four different movies were formed, assuming frames were formed by **(i)** incoherent imaging, **(ii)** coherent darkfield imaging (no reference field), **(iii)** coherent interference imaging, where the reference field has the same amplitude as each emitter PSF’s peak amplitude, and **(iv)** coherent interference imaging, where the amplitude of the reference field is 100 times greater than the peak amplitude of each emitter. For each movie, the mean intensity image and auto-cumulant images (*τ* = 0) up to the 8^th^-order are shown (top), together with the cross-section of the line passing through both emitters (bottom). Isotropic rotational diffusion was performed with a time step Δ*t* = 0.01*D*_*r*_, where *D*_*r*_ is the rotational diffusion coefficient. Movies were composed of 100,000 frames and simulated by fixing the two emitters *r*_0_/2 apart (blue dots), where *r*_0_is the Airy disk radius of the PSF for incoherent imaging.

Another observation that can be made from Fig. 1 is that for both incoherent imaging (Fig. 1i) and interference imaging with a significantly strong reference field (Fig. 1iv), the SOFI images exhibit two super-resolved spots in some orders, but not others. In general, both of these modalities support the application of SOFI algorithms since their resulting images do not include significant cross-terms between the PSFs of neighboring emitters/scatterers. The origin of these artifacts in certain orders can be explained by calculating the cumulants from the expected intensity fluctuations. For the case of particles that undergo isotropic rotational diffusion and a polarization-sensitive imaging modality such as fluorescence anisotropy imaging or interference imaging with a strong reference field, one can model the stochastic emitted/scattered light intensity from each particle as a random walk on a sphere, with the time-dependent intensity being proportional to the projection on one of the sphere’s axes. Under this assumption, one can derive the n^th^ moment *G*_*n*_ for the stochastic intensity trajectory of each emitter/scatterer:

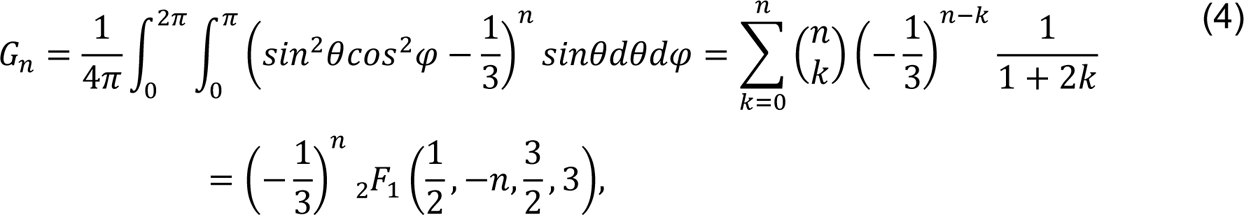

Where 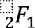 is the hypergeometric function. The corresponding cumulants can be calculated using the recursive formula described in Ref. [45], and are shown in Fig. 2. One notes the zero-crossing of the cumulant value at the 6^th^-order, resulting in SOFI images that do not display two resolved spots corresponding to the two point sources, as demonstrated for the case of incoherent imaging (Fig. 1i) and interference imaging with strong reference field (Fig. 1iv).

**Fig. 2.**
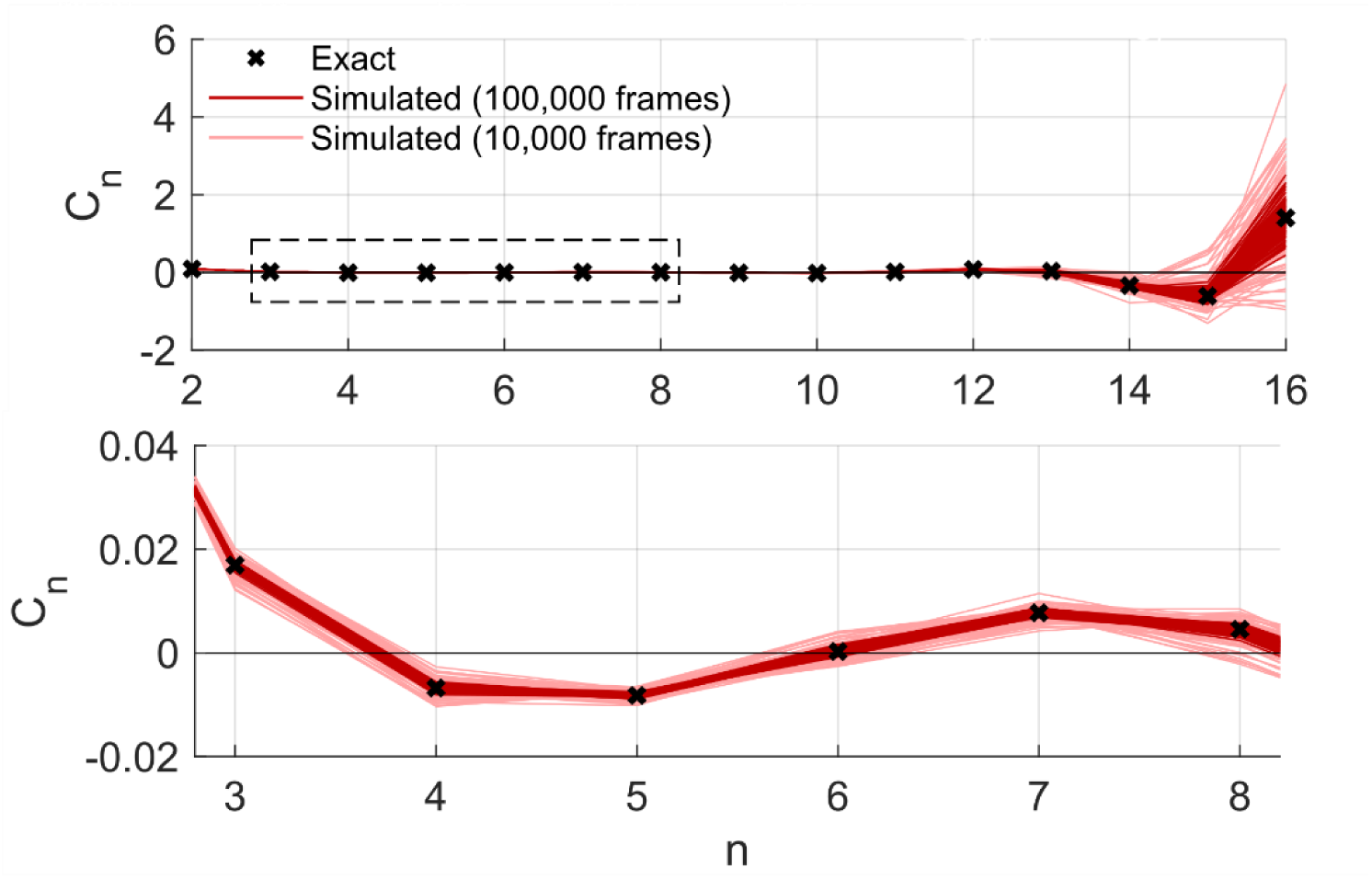
High-order cumulant analysis of rotationally diffusing emitters/scatterers. Simulated intensity trajectories of rotating scatterers were produced by assuming an isotropic random walk on a sphere. In order to emulate the anisotropic scattering of a nanorod with one main scattering axis, the intensity at each time point of the random walk was defined as the projection of the position on the sphere onto one of its axes. Auto-cumulants *C*_*n*_(*τ* = 0) values of simulated trajectories were calculated up to the 16^th^-order (top panel), assuming a trajectory composed of 100,000 (red) and 10,000 (light red) time points. The bottom panel shows a zoomed-in view of the orders marked by a dashed square. The theoretical cumulant values, calculated using Eq. 4, are shown for each order (black cross). Due to the zero-crossing of the cumulant value at the 6^th^-order, the SOFI algorithm does not resolve the two point emitters in the 6^th^-order SOFI image in Fig. 1(i) and 1(iv).

Cases in which the cumulant value approaches zero are particularly susceptible to cusp artifacts [46], occurring whenever two neighboring emitters/scatterers form cumulants of opposite signs. As shown in Fig. 2, the more statistically limited the trajectory duration used to calculate each cumulant, the greater the distribution of its values for a given diffusion process. For a wide enough cumulant value distribution, and to a greater extent in cases where the theoretical cumulant value approaches zero, the cumulant of two temporally fluctuating emitters/scatterers can be of opposite sign, even when created by identical diffusion processes. Fig. S4 shows an example of SOFI analysis of the interference-based movie simulated in Fig. 1, where only 10% of the movie duration is processed. One notes that the resulting SOFI images of different orders are highly variable between the various portions of the movie assumed for the analysis due to the high variability of cumulant values produced from each scatterer. This variability is caused by several factors. In the ideal case where the cumulant value from each scatterer is large compared to the cross terms, and both are of the same sign, the SOFI image results in two spots that become better resolved as the SOFI order increases. If only one of the two scatterers produces a large cumulant value, it will contribute more significantly to the corresponding SOFI image. In the case of large cumulant values of opposite signs, one observes a cusp artifact between the two scatterers. In cases where the cumulant value from each scatterer is significantly smaller compared to the cross terms, the resulting SOFI image is comprised mainly of the cross term, appearing as a spot located between the two scatterers. However, if the cumulant values from the two scatterers are comparable to the cross terms, a complex pattern emerges, where, for example, one observes three resolved spots. The SOFI images corresponding to the statistically limited movies, shown in Fig. S4, demonstrate the different cases. We note that the optimal number of frames for artifact-free SOFI also depends on the image acquisition rate. In general, the acquisition rate has to be rapid enough to capture the fluctuating optical signal from each individual particle in a manner that will produce non-negligible correlations between consecutive frames. However, temporally oversampling these intensity fluctuations will result in an unnecessarily large number of acquired frames. For the simulations described here, we sampled the rotational diffusion process with a time interval of Δ*t* = 0.01*D*_*r*_, where *D*_*r*_ is the rotational diffusion coefficient. Other fluctuative systems will require different optimal acquisition rates and total number of frames, which should be optimized for the particular underlying stochastic process in the imaged sample.

An additional important consideration that may affect coherent scattering-based imaging is the contribution of electromagnetic coupling between neighboring scatterers. It is likely that in cases where such coupling is significant, a correlation between otherwise independently fluctuating scatterers emerges. In order to assess the significance of electromagnetic coupling in practical applications, we produced simulated movies, via full wave-optical calculations [47, 48], composed of images formed by two rotating neighboring dipoles, taking into account the dipole-dipole interaction. This interaction is described by the following equations:

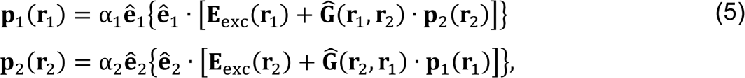

where **p**_1_(**r**_1_) and **p**_2_(**r**_2_) are the electric dipole moments with orientation unit vectors **ê**_1_ =**p**_1_/|**p**_1_| and **ê**_2_ = **p**_2_/|**p**_2_| that are induced in the first and second particle at positions **r**_1_ and **r**_2_ Here, E_exc_(**r**) is the external exciting electric field, and **Ĝ**(**r**_2_, **r**_1_) is a tensor describing the electric field generated at position **r**_2_ by a unit dipole at position **r**_1_. The constants α_1_ and α_2_ are the polarizabilities of the two particles, which generally depend on their size and shape. The solution of these coupled equations for **p**_1_(**r**_1_) and **p**_2_(**r**_2_) is given, in compact vector-matrix notation, by

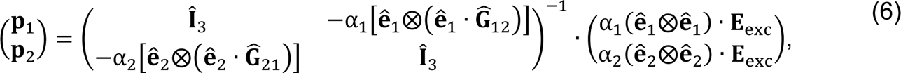

where the six-dimensional column vector on the l.h.s has the first three components **p**_1_and the last three components **p**_2_, and the matrix on the r.h.s. is a block matrix of four 3x3 matrices as shown, with Î_3_ denoting a 3x3 identity matrix, and the tensor product ⨂ between two vectors defined in the usual way by (a⨂b)_*jk*_ = a_*j*_b_*k*_.

Fig. 3 shows the resulting SOFI images for simulated interference imaging movies with the two neighboring dipoles rotating in a manner identical to the ones assumed for Fig. 1. Figs. 3(i) and 3(ii) show the resulting SOFI images for the case without and with dipole-dipole interaction, respectively, where the polarizabilities *α* of both rotating dipoles in the latter case were taken to be as one calculated for gold nanoparticles 40 nm in diameter [49]. One observes that under these assumptions, the effect of electromagnetic coupling between the dipoles is negligible, apart from slight influence in the 6^th^ SOFI order, where, as described above, the contribution of the cross-term between the two intensity trajectories of both particles is already very significant. We note that while both Fig. 1(iv) and Fig. 3(i) describe similar conditions for interference imaging of two non-interacting particles, the latter contains the added complexity of the angular dependence in the formed PSF from each dipole. Thus, one can notice slight differences between the SOFI images in Figs. 1(iv) and 3(i). These results demonstrate that dipole-dipole interaction can be likely neglected in many practical cases of scattering-based super-resolution imaging. However, for very high polarizabilities, or very close particles, such interactions will eventually lead to more significant effects that hinder super-resolution imaging. In order to demonstrate this point, we increased the polarizabilities of both particles by a factor of 10 and 100, shown in Figs. 3(iii) and 3(iv), respectively. There, one notices a greater difference from the non-interacting case shown in Fig. 3(i). In the extreme case of very high polarizabilities, as in Fig. 3(iv), dipole-dipole coupling removes the statistical independence between the stochastic scattered intensity trajectories of both particles, resulting in significant contributions from the cross-cumulant between neighboring particles. Thus, the corresponding SOFI images in Fig. 3(iv) contain only one spot located halfway between both dipoles. We note that the above electromagnetic simulations approximated both particles as point dipoles. However, particle geometry can play a significant role in image formation in cases where the particles’ size is comparable with the distance between them. We note that the considerations described in this section for a system comprising of two point particles can be extended to multi-particle systems.

**Fig. 3.**
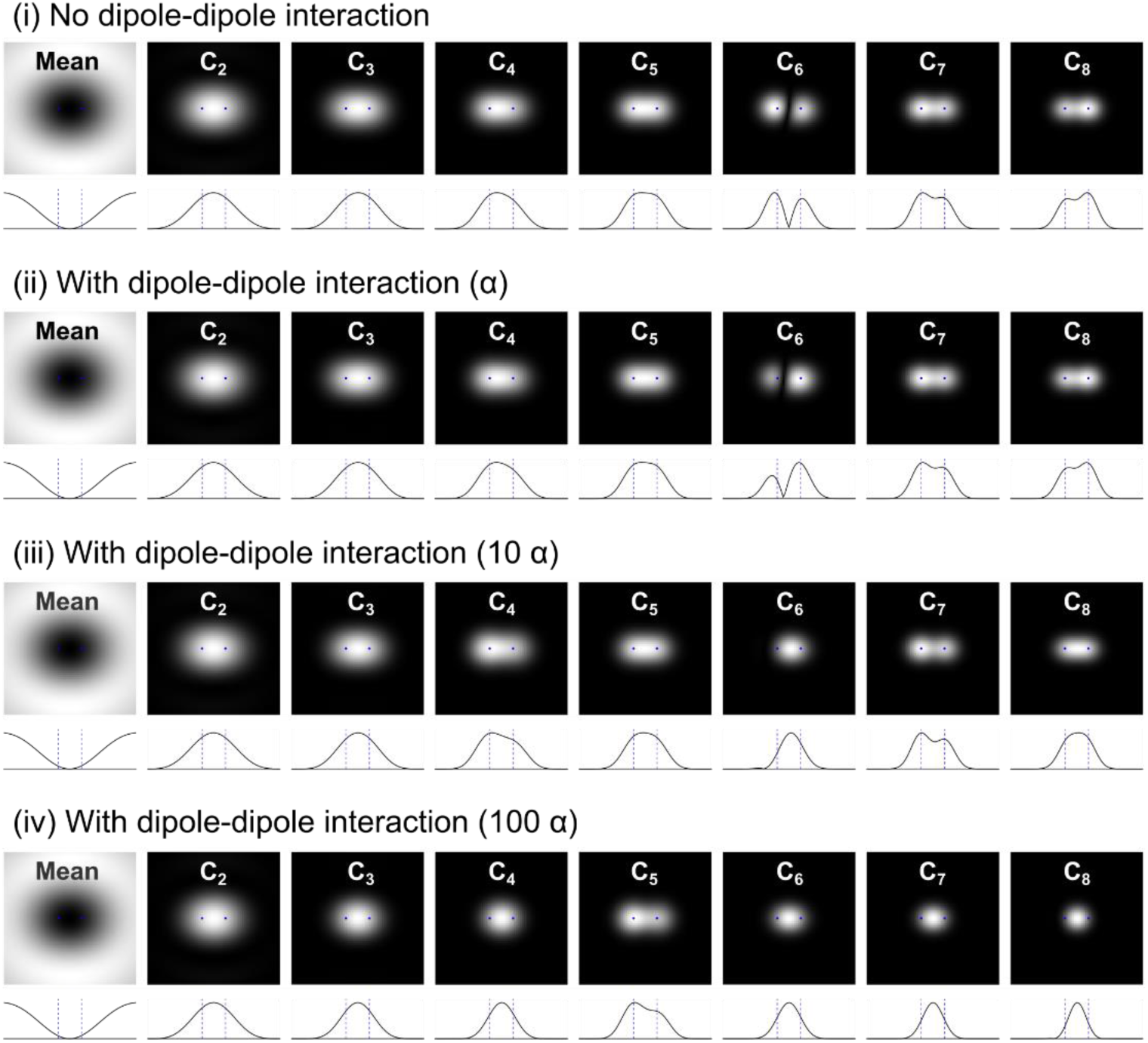
The effect of dipole-dipole interactions on scattering-based SOFI. Simulated interference imaging movies of two neighboring rotationally diffusing dipoles were produced assuming **(i)** no dipole-dipole interaction, and **(ii)** dipole-dipole interaction calculated for point dipoles with polarizabilities associated with gold nanoparticles that are 40 nm in diameter. For each movie, the mean intensity image and auto-cumulant images (*τ* = 0) up to the 8^th^-order are shown (top), together with the cross-section of the line passing through both emitters (bottom). Movies composed of 100,000 frames were produced by modeling an ideal interference imaging system with a strong reference field (*E*_*ref*_ = 100*E*_*scat*_) operating at a wavelength of 640 nm, with the two dipoles spatially fixed 160 nm apart in a water environment above a glass substrate, generating far-field images captured via an imaging system with 100X magnification, a numerical aperture of 1.2 and an effective pixel size of 13.5 nm. The rotational diffusion trajectory of each dipole was taken to be identical to the one used for generating the movies shown in Fig. 1. In order to accentuate the effect of dipole-dipole interactions, **(iii)** and **(iv)** show the resulting SOFI images calculated for movies produced by assuming polarizabilities that are 10x and 100x higher, respectively, than the one used for case **(ii)**.

Finally, we show in Fig. S5 the theoretical prediction for SOFI analysis performed on incoherent and interference-based coherent imaging of one particle translocating between two neighboring positions. This example also illustrates the case of two neighboring fluctuating emitters/scatterers in a completely anti-correlated manner. Due to the statistical dependence between the two fluctuating emitters/scatterers, the cross-term between the two particles does not vanish, resulting in its significant contribution to the resulting SOFI images. For this case, the resulting SOFI images of all orders, for both the incoherent and interference-based coherent imaging modalities, show two spots centered further apart than the two translocation positions.

### B. Experimental Demonstration of Scattering-based SOFI

In order to demonstrate the application of SOFI algorithms to scattering-based imaging modalities, we constructed a custom-built microscope capable of simultaneous reflection interference and epifluorescence imaging, as illustrated in Fig. S2 and further described in the ‘Materials and Methods’ section. Scanning electron microscopy (SEM) was used to observe the nanoparticles that produced the scattered light. Correlative imaging between optical and SEM microscopies was performed using an indexed chrome micro-grid fabricated on top of the glass coverslip (Fig. S3). Fig. 4A shows an example of such a dual imaging modality experiment, where two individual AuNRs, as seen from the SEM image in Fig. 4B, form a destructive interference pattern in the interference image.

**Fig. 4.**
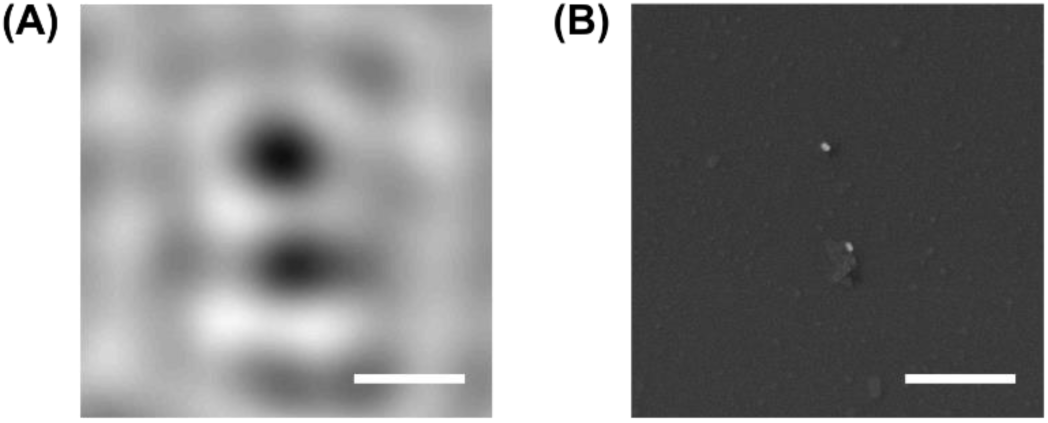
Correlative imaging: Interference-based and SEM imaging of AuNRs. **(A)** Two AuNR monomers were imaged via reflection interference microscopy, each forming a destructive interference pattern. Interference microscopy was performed with a 640 nm incoming laser beam. Background correction was applied to the raw interference image as described in the ‘Materials and Methods’ section. **(B)** SEM image of the same region shown in (A), localized using a grid pattern fabricated on the glass coverslip. Scale bars: 500 nm.

In a manner analogous to nanoscale calibration samples used for fluorescence-based super-resolution microscopies [41, 50], we formed scattering-based nanorulers by binding AuNRs at two defined positions on rectangular DNA origami structures. As illustrated in Fig. 5(A), the two AuNRs bind at the opposing corners of the rectangular origami structure. In addition, four fluorophores (ATTO 647N) were bound at the center of the structures, serving as a verification that the scattered light from the AuNRs is associated with a DNA origami structure. Figs. 5B and 5C show AFM images of the origami structures and SEM images of the bound AuNRs, respectively. Such scattering-based nanostructures can be utilized to test and develop scattering-based super-resolution methodologies. In order to test the feasibility of sSOFI, we acquired reflectance interference movies capturing fluctuations of the scattered light created by the rotational diffusion of these bound AuNRs, which scatter light in an anisotropic manner. Fig. 6A and 6B show the mean interference and SEM image of a region containing two individual AuNRs, and one AuNR dimer. We note that the two AuNRs in the dimer seem to be in contact with one another in the SEM image (Fig. 6B). We observed such contact between AuNRs in the SEM images of several AuNR dimers, verified to be associated with the DNA origami structure by the colocalized fluorescence signal from the fluorophores bound to the structure. This contact between the AuNRs may have occurred due to a drying artifact during sample preparation for SEM imaging, which included an evaporation process prior to the sputtering of the conductive iridium layer. Thus, it is likely that the two AuNRs in such dimers were not in contact with each other prior to evaporation, and were able to rotationally diffuse. Since the interference imaging detection path described in this work was polarization-sensitive, the rotational diffusion of these AuNRs was translated to intensity fluctuations of the detected scattered light. In the case of the dimer shown in Fig. 6, the combined PSFs demonstrated such temporally fluctuative patterns, which lent itself to further SOFI analysis. Fig. 6C shows the result of the SOFI analysis applied to an interference imaging movie, composed of 10,000 frames, of that sample region. We note that the number of captured frames was limited by the axial drift of the sample from the objective, which could significantly affect the resulting interference image for longer movies. In order to acquire a larger number of frames, one could incorporate a drift compensation mechanism into the imaging system. For the following demonstrations, SOFI analysis was performed by calculating the auto-cumulant value of each pixel at a time lag of *τ* = 1 frame, as it reduces the noise on the final SOFI image caused by any noise source that has the property of having no correlation between frames [24]. Speckle background correction was performed as described in the ‘Materials and Methods’ section. However, no further image processing, such as deconvolution, was utilized for the calculation of the SOFI images. In the SOFI-analyzed images shown in Fig. 6C, one observes two resolved spots exhibiting cusp artifacts in the odd SOFI orders and one spot in the even SOFI orders. For comparison, Fig. S6 shows the corresponding correlation function images from which the cumulant SOFI images were calculated. In these correlation function images, one observes that the AuNR dimer corresponds to primarily one spot in the high correlation orders. The resolvement of two spots in some SOFI orders in Fig. 6C, and its absence in the mere correlation function images shown in Fig. S6C, is the predicted outcome of the cumulant-based analysis utilized in SOFI, as it eliminates the contribution from cross-terms between neighboring particles that are present in the correlation function [24]. Two additional examples are shown in Fig. S7. There, two resolved spots, still exhibiting cusp artifacts, are present in even SOFI orders for the first example. The second example in Fig. S7 shows a more complicated pattern – while some SOFI orders contain two resolved spots, the orientation of the AuNR dimer is slightly different between some SOFI orders. These examples exhibit practical complexities that emerge in SOFI analysis in general [46]. However, since sSOFI can allow for both higher acquisition rates and longer acquisition durations as it is essentially free of photobleaching, the possibility of calculating SOFI images up to very high orders makes these complexities more apparent. From these examples, it is evident that the rotational diffusion of the AuNRs leading to intensity fluctuations does not necessarily follow the simple isotropic rotational diffusion model described in the previous section. Moreover, asymmetric or distorted PSFs introduce further complexities, especially in high orders, where asymmetries are further distorted as the effective PSFs in the SOFI image are powers of the original captured PSF. Such highly asymmetric neighboring PSFs can form complex patterns in the final SOFI image. While these factors introduce challenges to sSOFI, it is possible to address them experimentally and computationally. For example, it may be possible to create better-controlled fluctuating scattered light signals by implementing an approach similar to points accumulation for imaging in nanoscale topography (PAINT) [51] and DNA-PAINT [41], where scatterers are allowed to transiently bind to individual sites that can be resolved via SOFI. Utilizing an approach that offers reversible binding/unbinding kinetics, as in DNA-PAINT, can allow for further control and optimization of fluctuation kinetics. Moreover, since photobleaching is not an experimental concern in scattering-based imaging, one could develop fluctuating scattering probes by utilizing the opening/closing kinetics of DNA hairpins conjugated to scattering nanoparticles. In addition, asymmetric and distorted PSFs can be accounted for by deconvolving the interference images with known PSFs captured from static nano-scatterers scanned across the imaged sample area. Such approaches can enhance the capabilities of sSOFI and allow for the reduction of artifacts when calculating high cumulant orders. In addition, since the dimensions of the AuNRs in this demonstration are comparable to the distance between their binding sites on the DNA origami structure, a more detailed electrodynamic study of the scattering problem, accounting for nano-plasmonic effects, is required in order to describe the formed image [52, 53].

**Fig. 5.**
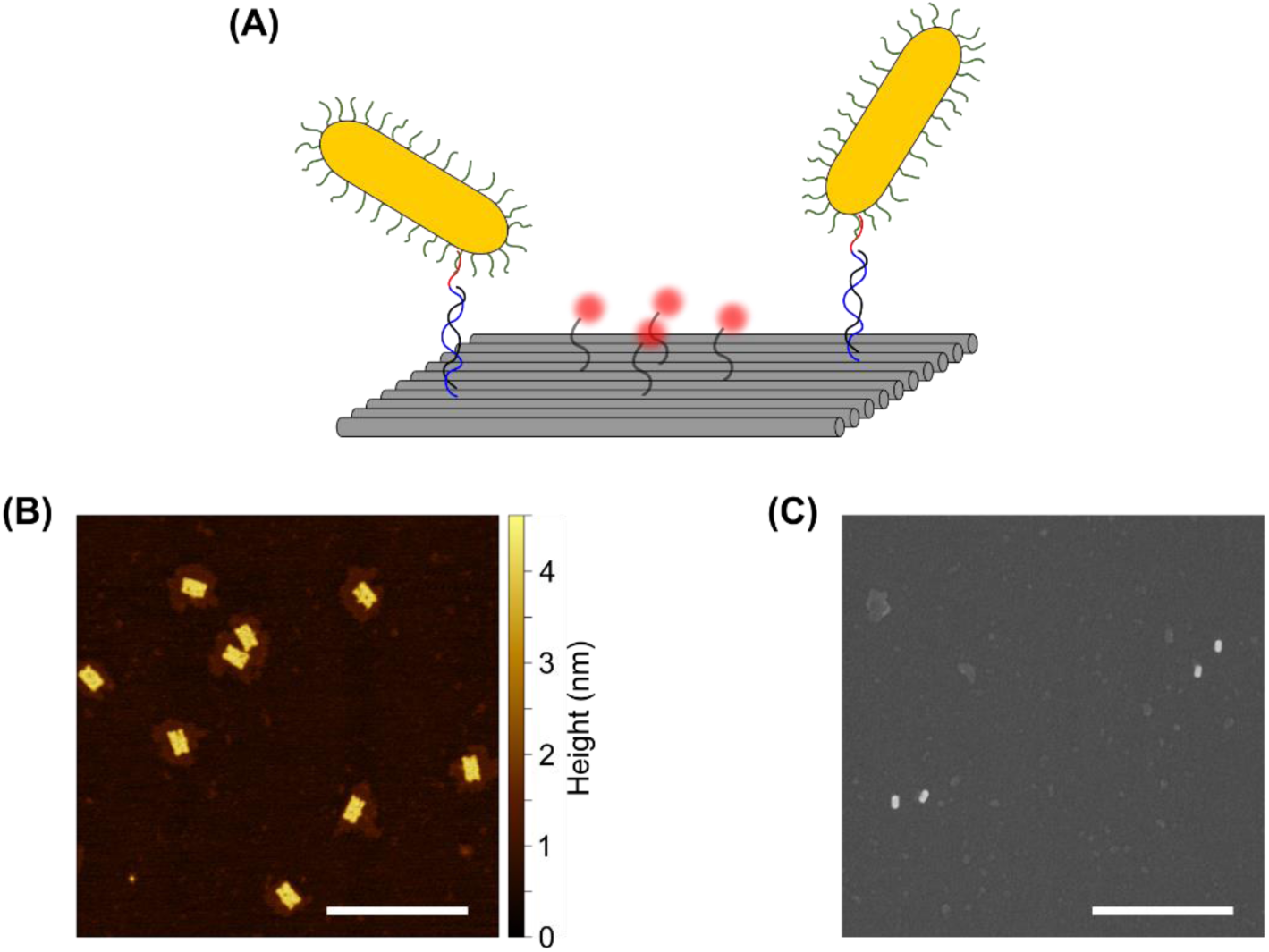
DNA origami-based gold nanorod dimers. **(A)** A scheme of the rectangular origami structure used in this work, binding two gold nanorods via two DNA strands ∼80 nm apart, and four fluorophores used for colocalization between the origami structures and gold nanorods. **(B)** AFM image of the folded rectangular origami structures. **(C)** SEM image of gold nanorods bound to the origami structures, forming dimer geometries. Scale bars: 500 nm.

**Fig. 6.**
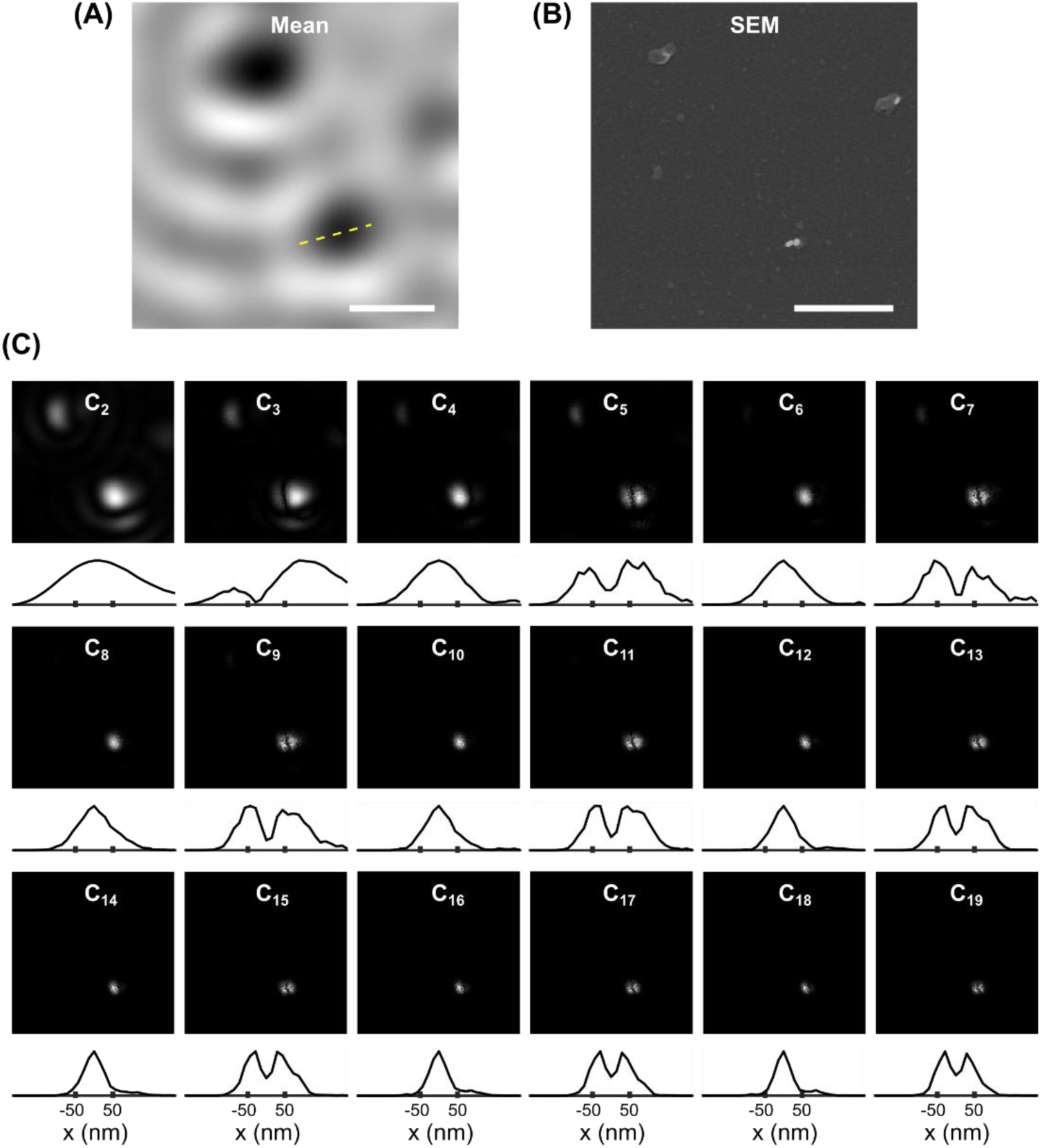
SOFI analysis on interference-based imaging of gold nanorod dimers. Reflectance interference microscopy and corresponding SOFI analysis of gold monomer and dimer nanorods. **(A)** and **(B)** show the averaged background-corrected iSCAT image and SEM image, respectively. A movie composed of 10,000 frames was acquired (exposure time = 1 ms) using a polarized incoming beam, which results in fluctuating PSFs for the case of rotationally diffusing nanorods. Image acquisition buffer consisted of 5 mM Tris-HCl, 10 mM MgCl_2_, 1 mM EDTA, and 0.05% (v/v) Tween 20 (pH 8). **(C)** shows the SOFI auto-cumulant images (*τ* = 1 frame) up to the 19^th^-order. For each order, the cross-section of the region indicated with a yellow dashed line in **(A)** is shown below each SOFI image. Scale bars: 500 nm.

### C. Scattering-based SOFI as a Fluctuation-based Novelty Filter

In this final section, we demonstrate the application of sSOFI as an imaging processing approach for highlighting dynamic processes captured by interference imaging. In addition to its use for super-resolution imaging, sSOFI can be utilized as a novelty filter and provide a quantitative measure for cellular activity that results in fluctuating scattering from biomolecules, in a manner similar to the different variants of imaging-based correlation spectroscopy [54, 55]. Fig. 7 shows the application of SOFI analysis to interference microscopy movies of membrane protrusions in live cells. Here, fluctuating scattered fields from biomolecules were analyzed via SOFI algorithms for contrast enhancement. Membrane labeling was used to identify two neighboring membrane protrusions (Figs. 7B-C), which produced very little contrast in both the brightfield (Fig. 7A) and interference images (Fig. 7D). One of the protrusions, due to the high degree of fluctuating scattered fields from the biomolecules diffusing and transporting within it, appears in the SOFI images as a bright structure on top of a dark background (Figs. 7E-F). Such diffusion and transport processes can produce fluctuating optical signals even from isotropic scatterers, and can be straightforwardly highlighted by SOFI algorithms. The structure displays noticeable spatial oscillations, which can be either due to morphological properties of the sample, such as focal contacts, or imaging-related, such as the background speckle pattern created by the scattering from the glass-water interface. The use of sSOFI as an activity-sensitive contrast mechanism can be applied simultaneously with conventional fluorescence imaging with labeled biomolecules, even with the same excitation light, providing information on the overall activity of biomolecules in the local environment without additional labeling.

**Fig. 7.**
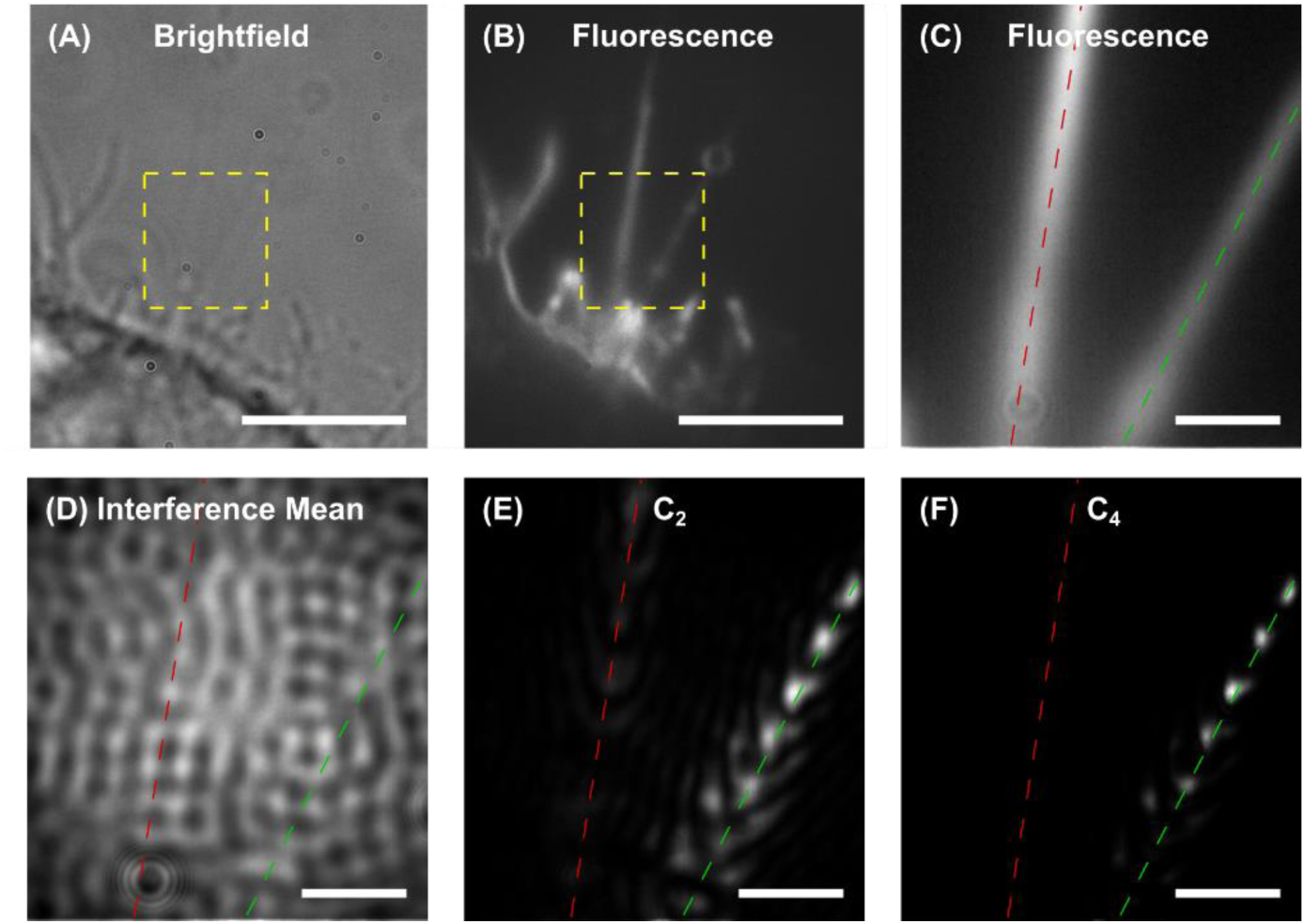
Scattering-based SOFI analysis of membrane protrusions exhibiting fluctuative scattering. Fluorescently labeled membrane protrusions of a HEK293 cell were imaged using **(A)** brightfield, **(B**-**C)** fluorescence, and **(D)** interference microscopies. The dashed yellow squares in **(A)** and **(B)** indicate the zoomed-in regions shown in **(C**-**F)**. The cell membrane, including its membrane protrusions, was labeled with Di-8-ANEPPS. For interference imaging, a movie composed of 1000 frames was acquired (exposure time = 1 ms) using a 470 nm incoming laser beam. Cellular activity that resulted in the translocation of biomolecules generated a fluctuating scattering signal, which was then highlighted by SOFI analysis. The dashed green and red lines indicate the active and inactive protrusions, respectively. **(E)** and **(F)** show the 2^nd^ and 4^th^-order SOFI auto-cumulant images (*τ* = 1 frame) for the raw interference intensity movie. Scale bars: **(A**-**B)**: 5 µm, **(C**-**F)**: 1 µm.

## 4. Conclusion

In summary, we investigated the application of SOFI algorithms to coherent scattering-based imaging modalities. We showed that while dark-field-based imaging modalities cannot be super-resolved via SOFI algorithms due to a cross-term between the two scatterers appearing in the image formed at the camera plane, interference-based imaging modalities are suited for further super-resolution processing via SOFI algorithms. Since SOFI algorithms are based on uncorrelated fluctuating signals from neighboring dipoles, a mechanism for creating such fluctuations is essential for achieving super-resolved scattering-based images. While fluorescence blinking provides a source of signal fluctuations for incoherent, fluorescence-based SOFI, scattering-based imaging requires an alternative mechanism. Here, we utilized the rotational diffusion of anisotropic nanoparticles that, when imaged in a polarization-sensitive modality, can provide a source of fluctuating scattered field from each nanoparticle. In order to demonstrate the application of SOFI for achieving super-resolved images from interference-based movies, we formed AuNRs dimers by tethering them to DNA origami constructs. Such engineered scattering nano-rulers can provide a platform for the further development of scattering-based super-resolution approaches. In addition, by imaging membrane protrusions, we demonstrated the use of sSOFI as a novelty filter that can be utilized in order to highlight dynamic processes that produce fluctuating scattering fields. This label-free contrast-generating image processing approach could be applied to visualize dynamic events and processes captured in scattering-based imaging.

The development of scattering-based super-resolution techniques can allow one to probe biological systems without the disadvantages of fluorescence imaging, which are inherently limited due to a finite photon budget. Practically, the development of sSOFI will require one to either optimize sample preparation in order to produce reliable fluctuative scattering, by, for example, rotational diffusion of anisotropic nanoparticles, or binding and unbinding kinetics, or rely on dynamic cellular processes such as diffusion and transport of small organelles or large macromolecules. Once achieved, such bleaching-free acquisition will allow for super-resolution imaging at an unprecedented temporal resolution and imaging duration.

## Data availability

All acquired and simulated movies used for the results presented in this work are available at Zenodo: https://doi.org/10.5281/zenodo.12612205. Home-written code (MATLAB) for the analysis shown in this work is available upon request.

