## Supplementary Figures for "Scattering-Based Super-Resolution Optical Fluctuation Imaging"

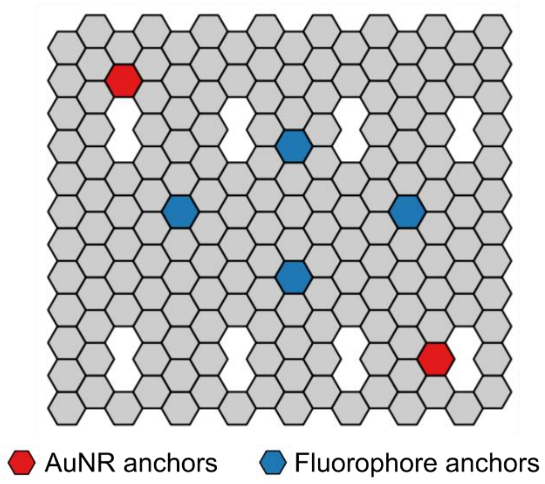


**Fig. S1. DNA origami binding strands positions.** The DNA origami structure was designed using the Picasso software package [1]. Dedicated binding strands were added to anchor two functionalized AuNRs approximately 80 nm apart (red tiles: Plate 1, C11 and Plate 2, F2), and four fluorophores (blue tiles: Plate 1, E7 and Plate 2, A5, A9, and E7), as further described in the 'Materials and Methods' section. Tile map, plate positions, and DNA sequences were obtained via the 'Design' module of the Picasso software.

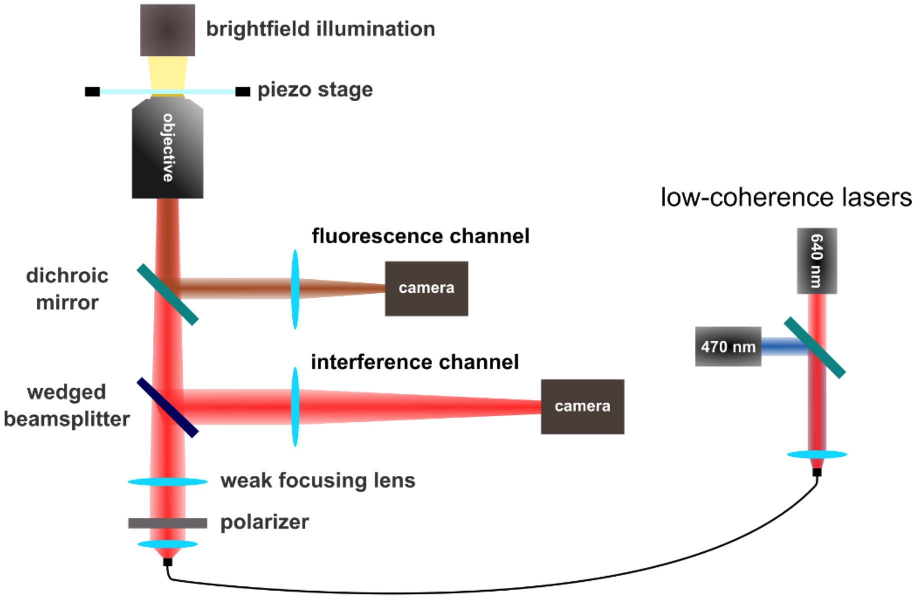


**Fig. S2. Simplified illustration of the optical scheme.** The optical scheme used for reflection interference and epifluorescence imaging modalities is further described in the 'Materials and Methods' section.

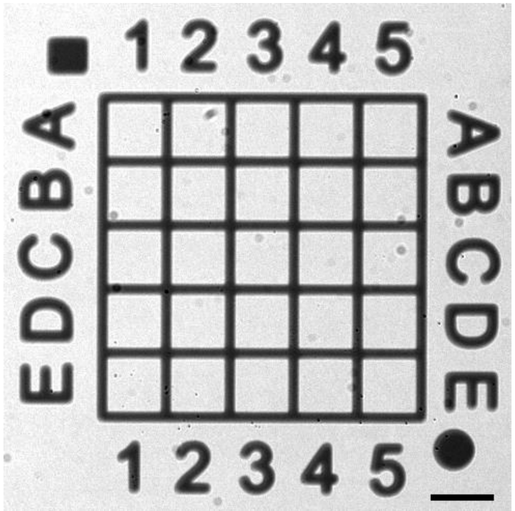


**Fig. S3. Micro-grid for optical-SEM colocalization.** In order to locate individual particles via both optical and SEM imaging, indexed grids were fabricated on top of glass coverslips. Each grid comprised an array of 20 µm x 20 µm cells fabricated photolithographically. Briefly, PMGI SF5 and AZ 1518 photoresist (MicroChemicals) were spin-coated on glass coverslips (Paul Marienfeld GmbH). The substrate was exposed to the grid pattern via a maskless lithography system (MLA 150, Heidelberg Instruments). Then, a 5 nm chrome layer was sputtered (Bestec GmbH) on the developed substrate. Finally, substrates were incubated in NMP for photoresist removal. Scale bar: 20 µm.

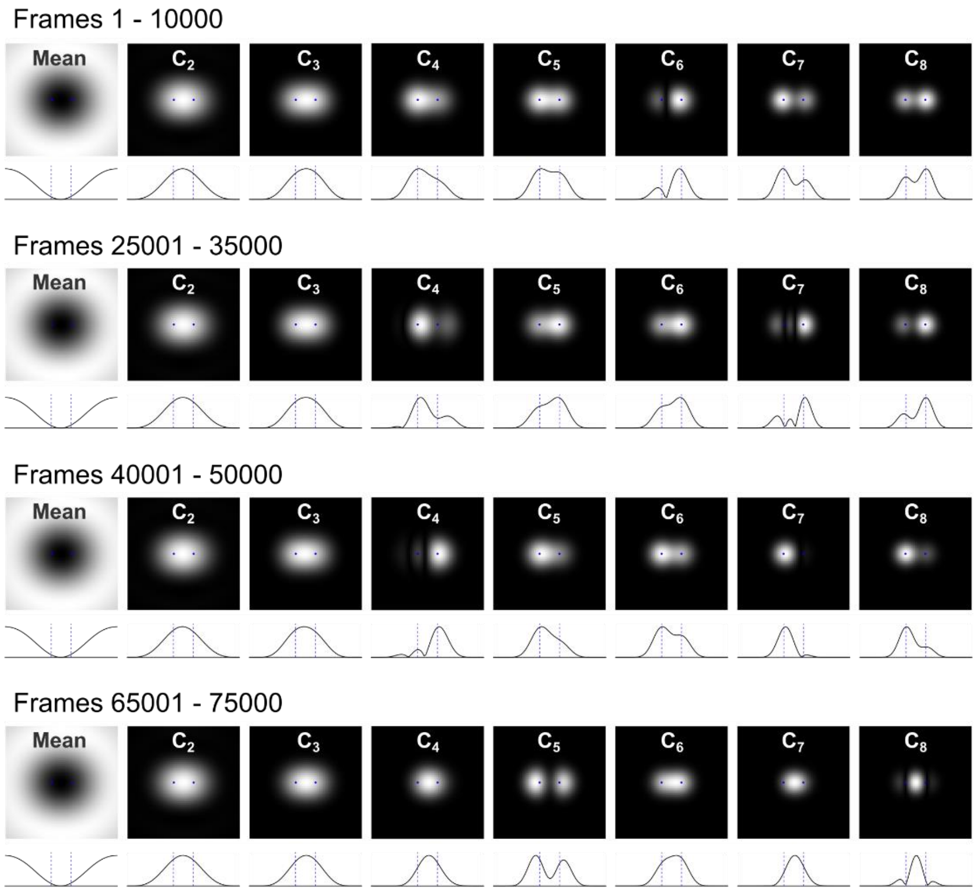


**Fig. S4. SOFI analysis of simulated interference imaging movies with limited statistics.** 10,000-frame sections of the simulated movie produced for Fig. 1(iv), which assumed two neighboring rotationally diffusing scatterers imaged via coherent interference imaging ($E_{ref} = 100E_{scat}$), were used for SOFI analysis instead of the total 100,000 frames. For each section, the mean intensity image and auto-cumulant images ($\tau = 0$) up to the 8$^{th}$-order are shown (top), together with the cross-section of the line passing through both scatterers (bottom). In such statistically limited datasets, the cumulants resulting from the fluctuating signals from each scatterer can significantly vary even when the signals from both scatterers follow the same diffusion process. As shown here, SOFI images calculated from such statistically limited movies can contain both cusp artifacts in cases where the cumulants of the two scatterers are of opposite signs, and a significant contribution from the cross-term between the two scatterers when the cumulants of the two scatterers are small or comparable to the cross-term, which forms an effective emitter located between the two real scatterers.

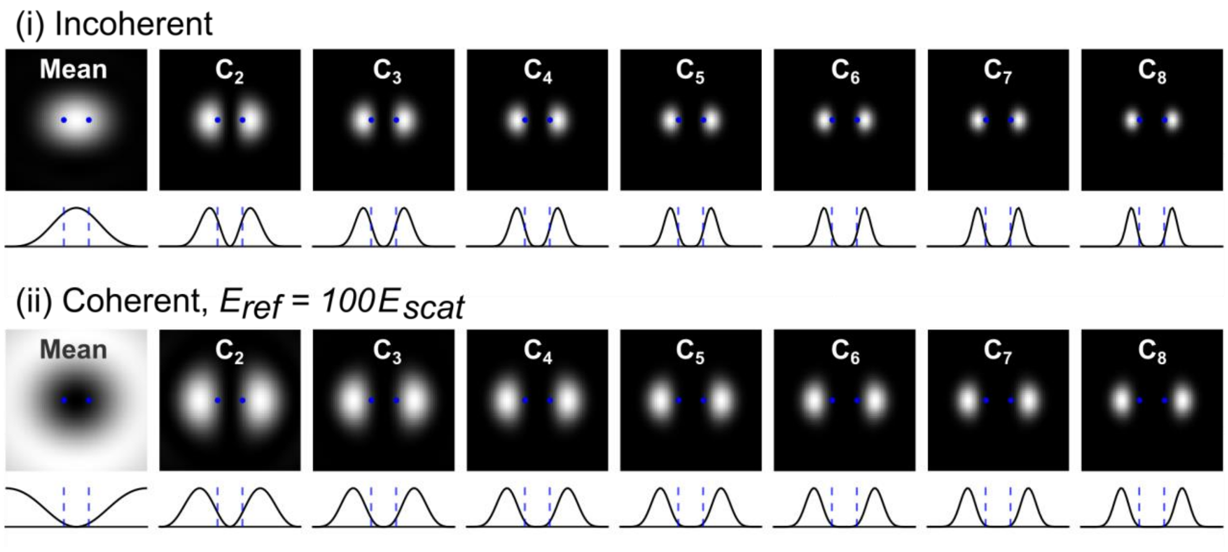


**Fig. S5. Cusp artifacts in high-order incoherent SOFI of a translocating scatterer.** Simulated movies and the corresponding SOFI images of an individual emitter translocating between two neighboring points in a Poissonian manner. Two different movies were formed for the same translocation trajectory, assuming frames were formed by **(i)** incoherent imaging and **(ii)** coherent interference imaging, where the amplitude of the reference field is 100 times greater than the peak amplitude of each emitter. For each movie, the mean intensity image and auto-cumulant images ($\tau = 0$) up to the 8$^{th}$-order are presented (top), together with the cross-section of the line passing through both emitters (bottom). Movies were composed of 100,000 frames and simulated by forming an emitter described by an Airy function that is translocating between two positions $r_0/2$ apart (blue dots), where $r_0$ is the Airy disk radius of the PSF for incoherent imaging. While the simulation assumed equal probabilities for the emitter being positioned in either of the two possible positions, it does not affect the SOFI analysis images in the case of noiseless imaging, as the fluctuation associated with the removal of a particle is identical to the fluctuation associated with the addition of one.

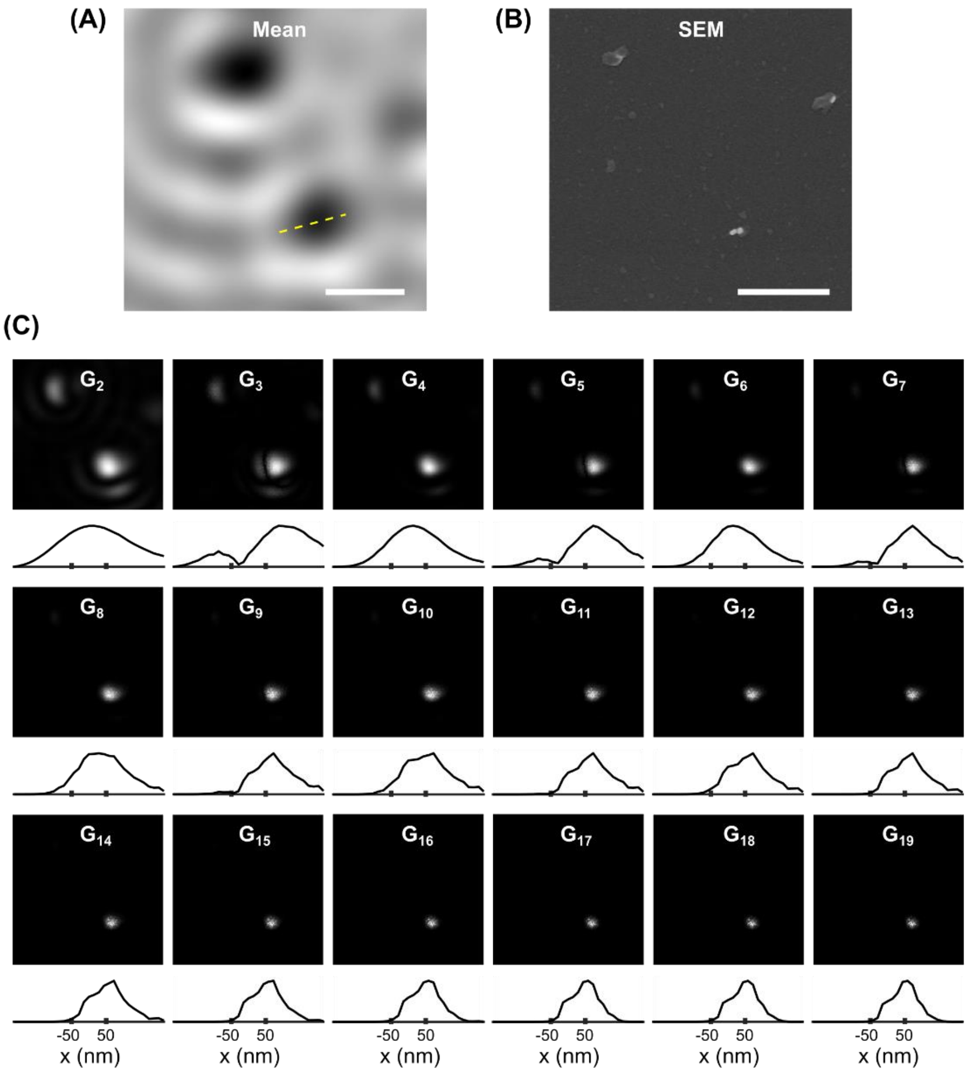


**Fig. S6. Correlation function analysis on interference-based imaging of gold nanorod dimers.** Autocorrelation image analysis of the gold monomer and dimer nanorods shown in Fig. 6. **(A)** and **(B)** show the averaged background-corrected iSCAT image and SEM image, respectively. A movie composed of 10,000 frames was acquired (exposure time = 1 ms) using a polarized incoming beam, which results in fluctuating PSFs for the case of rotationally diffusing nanorods. **(C)** shows the autocorrelation function images ($\tau = 1$ frame) up to the $19^{th}$-order, from which the auto-cumulant SOFI images shown in Fig. 6C were calculated. For each order, the cross-section of the region indicated with a yellow dashed line in **(A)** is shown below each autocorrelation image. As expected, the nanorod dimer does not produce a double-peaked image in the high-order autocorrelation images, in contrast to the high-order auto-cumulant SOFI images shown in Fig. 6C. Scale bars: 500 nm.

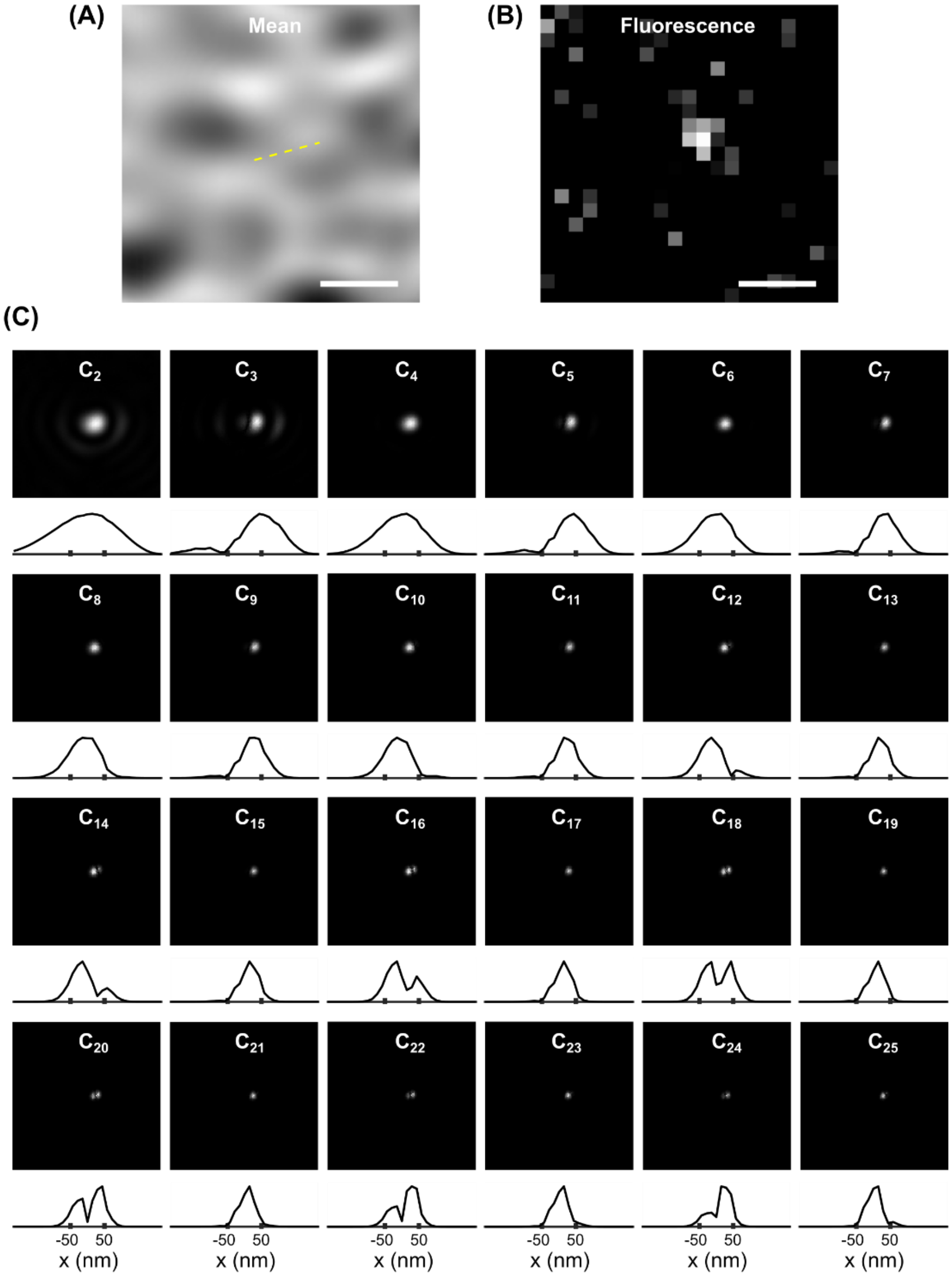

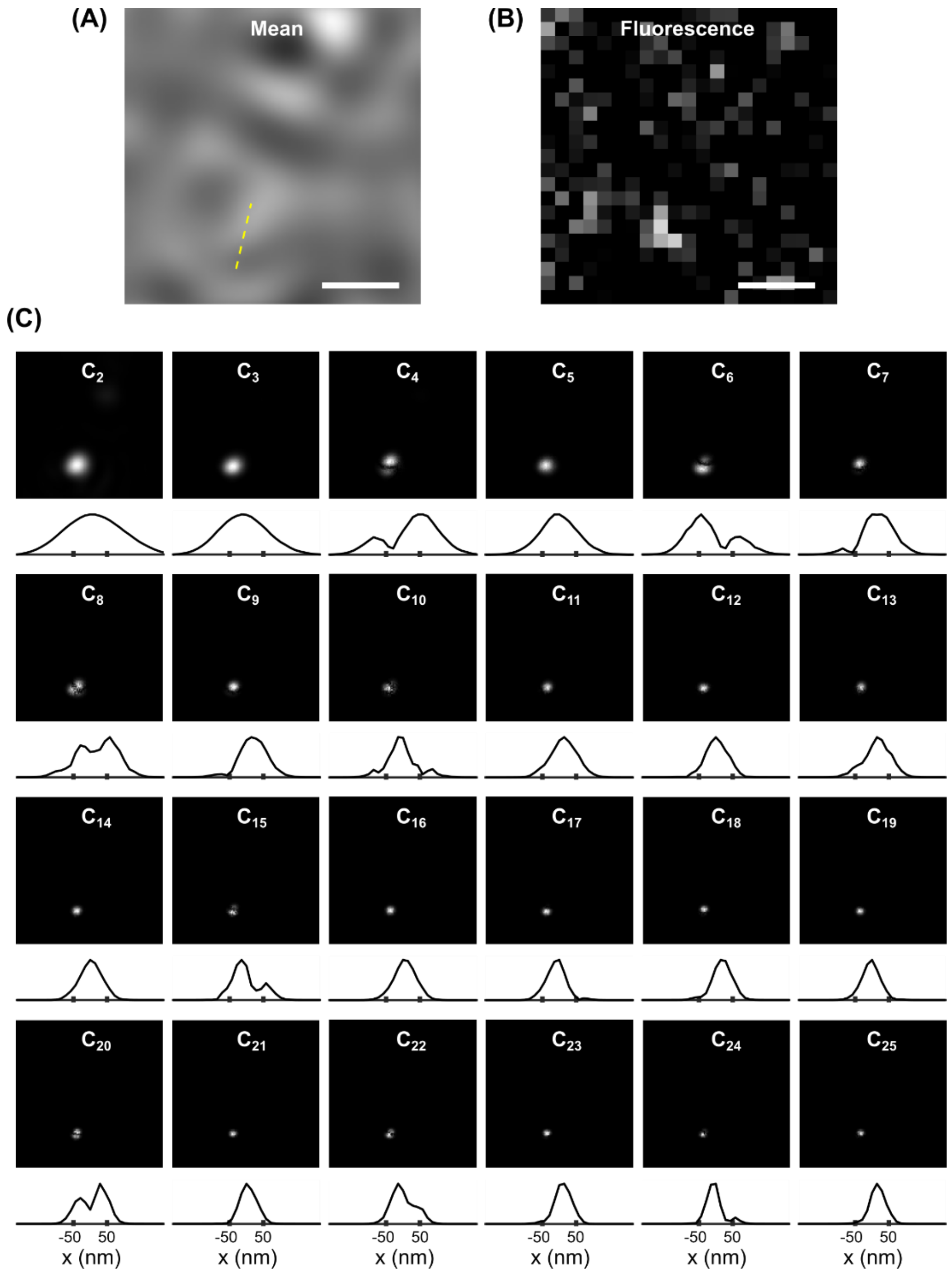


**Fig. S7. SOFI analysis on interference-based imaging of gold nanorod dimers.** Reflectance interference microscopy and corresponding SOFI analysis of gold monomer and dimer nanorods. **(A)** and **(B)** show the averaged background-corrected iSCAT image and fluorescence image of the DNA-tethered ATTO647 fluorophore, respectively. While SEM images of the optically imaged regions were not acquired, the colocalization between the scattered light from the AuNRs in the interference image and the fluorescence from the labeled DNA origami structures indicates that the fluctuating constructs were likely AuNR dimers. A movie composed of 10,000 frames was acquired (exposure time = 1 ms) using a polarized incoming beam, which results in fluctuating PSFs for the case of rotationally diffusing nanorods. Image acquisition buffer consisted of an aqueous buffer supplemented with 75% (w/w) glycerol containing 5 mM Tris-HCl, 10 mM $MgCl_2$, 1 mM EDTA, and 0.05% (v/v) Tween 20 (pH 8). **(C)** shows the SOFI auto-cumulant images ($\tau = 1$ frame) up to the 25$^{th}$-order. For each order, the cross-section of the region indicated with a yellow dashed line in **(A)** is shown below each SOFI image. Scale bars: 500 nm.
